# A Reporter Mouse for *In Vivo* Detection of DNA Damage in Embryonic Germ Cells

**DOI:** 10.1101/2020.01.29.925487

**Authors:** Jordana C. Bloom, John C. Schimenti

**Author notes:** Correspondence to: John C. Schimenti, Department of Biomedical Sciences, Cornell University, Ithaca, New York, 14853.

## Abstract

Maintaining genome integrity in the germline is essential for survival and propagation of a species. In both mouse and human, germ cells originate during fetal development and are hypersensitive to both endogenous and exogenous DNA damaging agents. Currently, mechanistic understanding of how primordial germ cells respond to DNA damage is limited in part by the tools available to study these cells. We developed a mouse transgenic reporter strain expressing a 53BP1-mCherry fusion protein under the control of the Oct4ΔPE embryonic germ cell-specific promoter. This reporter binds sites of DNA double strand breaks (DSBs) on chromatin, forming foci. Using ionizing radiation as a DNA double strand break-inducing agent, we show that the transgenic reporter expresses specifically in the embryonic germ cells of both sexes and forms DNA damage induced foci in both a dose- and time-dependent manner. The dynamic time-sensitive and dose-sensitive DNA damage detection ability of this transgenic reporter, in combination with its specific expression in embryonic germ cells, makes it a versatile and valuable tool for increasing our understanding of DNA damage responses in these unique cells.

## Introduction

In most contexts, DNA double strand breaks (DSBs) are damage-induced lesions which can arise as a by-product of normal cellular processes or from exogenous damaging agents (Featherstone & Jackson, 1999). If not repaired correctly, DSBs can be mutagenic and lead to deletions, translocations, and other chromosomal rearrangements. If left unrepaired, they can trigger cell death (Cannan & Pederson, 2016). The most common method of detecting DSBs is by immunolabeling with an antibody against different markers of DNA damage, such as γH2AX (Podhorecka, Skladanowski, & Bozko, 2010). While frequently used, this method requires cell fixation and depends on antibody specificity and sensitivity. More recently, a number of labs have begun to use fluorescent reporters of p53 binding protein 1 (53BP1) to quantify DSBs in individual living cultured cells (Karanam, Kafri, Loewer, & Lahav, 2012; Miwa et al., 2013). A major advantage of this method is that fixation and subsequent immunohistochemistry are not required for DSB visualization. Forgoing fixation enables visualization of DNA repair dynamics over time.

Preserving genome integrity in germ cells is essential for successful transmission of hereditary information to future generations and survival of a species. In many organisms, including mice and humans, germ cells are first specified during embryonic development (Ginsburg, Snow, & McLaren, 1990; Gomperts, Wylie, & Heasman, 1994; Tam & Snow, 1981). These primordial germ cells (PGCs), arising from a small progenitor pool, migrate and proliferate to subsequently form the entire adult germline (Gomperts et al., 1994; Tam & Snow, 1981). Perturbations to PGC development and growth, including those affecting genomic integrity of PGCs, can lead to a severe reduction or complete loss of germ cells in sexually mature adults (Hamer & de Rooij, 2018). Notably, mutations in a number of genes involved in DNA repair cause depletion of PGCs and consequently a reduction of germ cells post-natally, while having very subtle effects in the soma (Agoulnik et al., 2002; Luo et al., 2014; Luo & Schimenti, 2015; Nadler & Braun, 2000). Furthermore, mouse PGCs are particularly hypersensitive to ionizing radiation (IR)-induced DSBs, but the basis for this sensitivity remains unknown (Heyer, MacAuley, Behrendtsen, & Werb, 2000), in part due to the limited accessibility and numbers of fetal germ cells. Improvement to our current understanding of the mechanisms responsible for hypersensitivity of these cells to exogenous and endogenous DNA damage would benefit substantially from new research-enabling experimental tools. Here, we describe a transgenic mouse model for *in vivo* DSB detection specifically in fetal germ cells, alleviating technical barriers for studying DNA damage and repair in this difficult-to-access embryonic cell population.

## Results

### Functional Validation of the Oct4ΔPE-53BP1mCherry Reporter Transgene

To better understand DSB response dynamics in embryonic germ cells, we a built a reporter construct in which the Oct4ΔPE embryonic germ cell promoter (Yoshimizu et al., 1999) drives an established DSB-sensing reporter (Figure 1A) (Dimitrova, Chen, Spector, & de Lange, 2008). This promoter fragment (which lacks a proximal enhancer element) has been well characterized for its ability to drive specific expression in primordial and later stage fetal germ cells (Sabour et al., 2011; Szabó, Hübner, Schöler, & Mann, 2002; Yeom et al., 1996). The reporter is composed of mCherry fused to the Tudor binding domain of the 53BP1 protein and functions by binding to the H4K20me2 chromatin mark which becomes exposed adjacent to DNA DSBs (Dimitrova et al., 2008). In the absence of DSBs, the reporter is expressed in the nucleus in a diffuse pattern, but once DSBs are induced, the reporter forms foci at the sites of damage.

**Figure 1.**
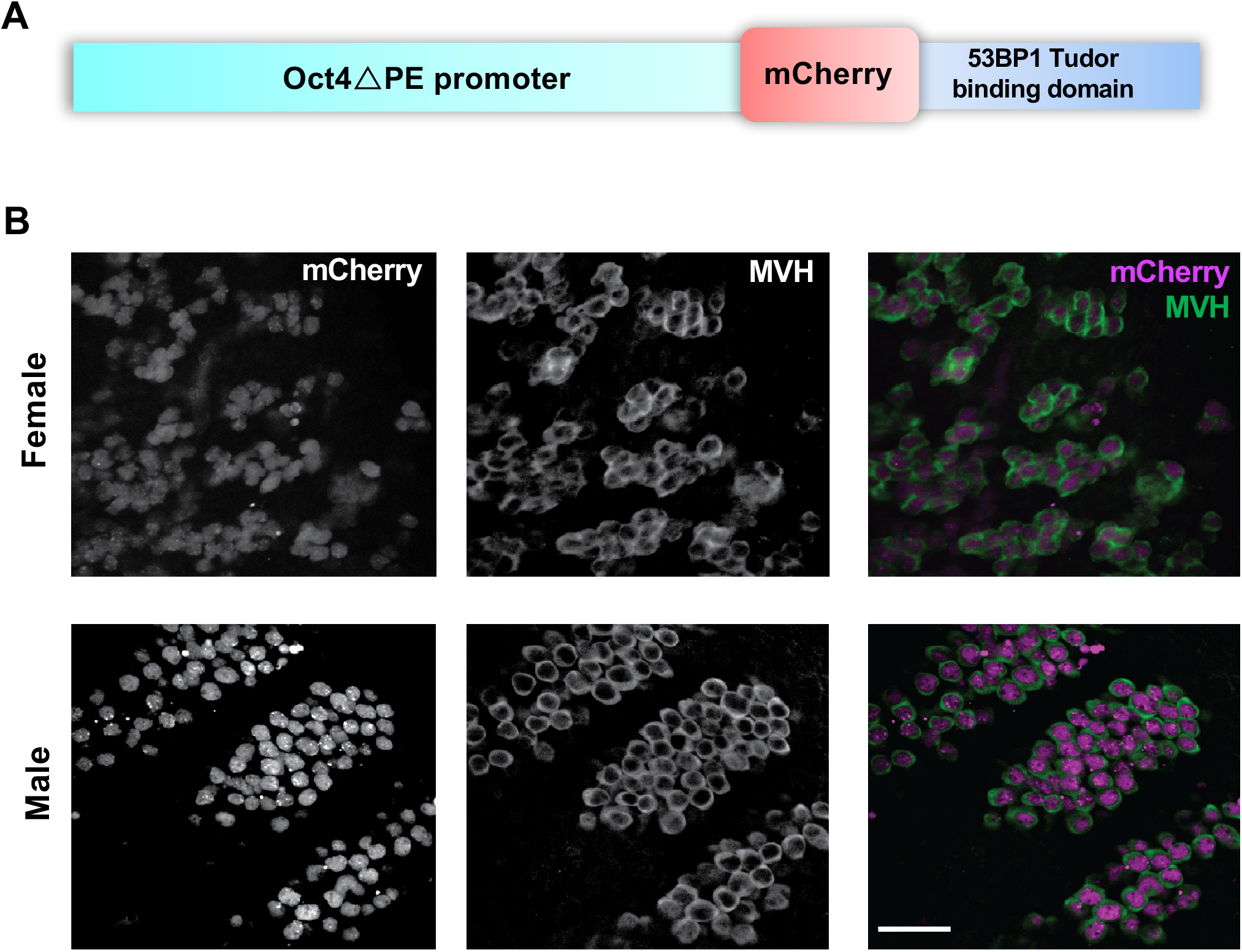
Creation of a mouse line expressing an Oct4ΔPE-53BP1-mCherry transgene specifically in male and female embryonic germ cells. (A) Schematic diagram of transgene construct with the Oct4ΔPE embryonic germ cell specific promoter (~20 kb) and the 53BP1-mCherry DNA double strand break sensing reporter (~2.2 kb). (B) Whole mount immunofluorescence images of E13.5 transgenic male and female genital ridges with panels on the left showing endogenous mCherry reporter localization, middle panels showing antibody labeling of the cytoplasmic germ cell marker MVH (DDX4), and panels on the right showing the corresponding merged image. Scale bar=50 μm

The Oct4ΔPE-53BP1mCherry reporter construct (Figure 1A) was microinjected into single cell mouse zygotes, and founder transgenics were identified via PCR with transgene-specific primers (see Table 1). Following germline transmission of the transgene, sexually mature transgene positive males and females were paired for timed matings. Gonads from male and female transgene positive embryos were collected at embryonic day 13.5 (E13.5), and co-labeled with the germ cell marker MVH (mouse vasa homolog; formally, DDX4) to assess if transgene expression was specific to germ cells in both sexes (Figure 1B). After confirming PGC-specific expression of the transgene, we validated that its expression does not have a detrimental impact on the fertility of transgene-positive adult female and male animals (Figure S1A-E).

We next sought to determine whether the DSB sensing reporter portion of the transgene was functional. To examine this, we exposed pregnant mice at 13.5dpc to a single low dose of IR (1Gy) and quantified focus formation 1 hour later compared to unirradiated transgene-positive counterparts (Figure 2A). Quantification of foci revealed that the reporter functions in both a dose-dependent (Figure 2B, C) and time-dependent manner with damage-induced foci being undetectable 24 hours after IR (Figure S2).

**Figure 2.**
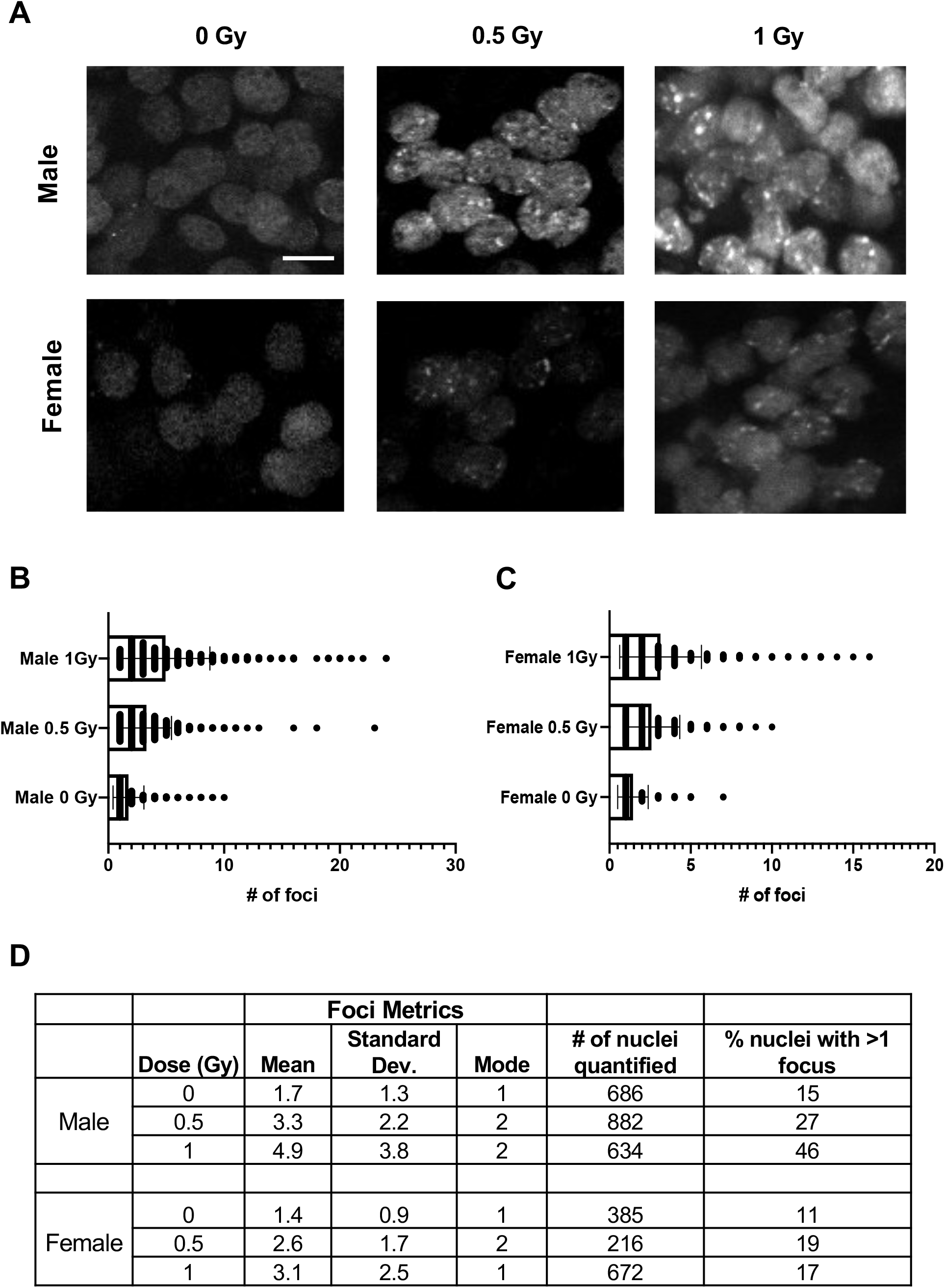
Primordial germ cells of Oct4ΔPE 53BP1-mCherry transgenic mice form radiation-induced foci in a dose-dependent manner. (A) Whole mount immunofluorescence images of E13.5 transgenic male and female genital ridges 1 hour after exposure to 0, 0.5, or 1 Gy of radiation. Scale bar=10 μM. (B) Foci quantification of transgenic male PGCs at the doses shown in A; p-values derived from unpaired Student’s t-tests where p<0.0001 for all comparisons shown. (C) Foci quantification of transgenic female PGCs at the doses shown in A; p-values derived from unpaired Student’s t-tests where p=0.002 for the 0.5 vs. 1 Gy statistical comparison and p<0.0001 for the 0 vs. 0.5 Gy and 0 vs. 1 Gy comparisons. (D) Table of reporter induction metrics in response to radiation including mean number of foci formed, mode, number of nuclei quantified per condition and the percentage of nuclei with more than one focus.

### Radiation-induced Reporter Foci Dynamics Differ Between Male and Female E13.5 PGCs

While IR cause both female and male PGCs to form mCherry foci in a dose-dependent manner, the average number of induced foci differed between the sexes (Figure 2B, C). At E13.5, female germ cells had fewer foci than males, and the percentage of nuclei with >1 focus in response to increasing IR doses remained fairly constant. In contrast, the percentage of nuclei with >1 focus steadily increased in male germ cells with increasing IR dosages (Figure 2D). We speculate that these differences might be due to the different developmental trajectories of the sexes at this time point. At E13.5, female germ cells begin to enter the leptotene stage of prophase I in meiosis I (Speed, 1982). Upon meiotic entry, the germ cells produce endogenously programmed DSBs that induce, and are required for, meiotic homologous recombination (HR) and pairing of homologous chromosomes (Handel & Schimenti, 2010). While repair of DSBs in meiosis occurs exclusively via HR pathway components, 53BP1 is associated primarily with the non-homologous end joining (NHEJ) DSB repair pathway. Therefore, its activity as a DSB responsive element may be down-regulated during early meiosis (Enguita-Marruedo et al., 2019), resulting in less 53BP1-based reporter foci detection in the female germ cells at this time.

### An Acute Dose of Radiation During Fetal Development Leads to a Reduction of Germ Cells in Males and Females at Birth

In order to assess the ramifications of DNA damage on PGCs, we performed the same experimental treatment conditions as previously described, but collected female and male gonads at birth rather than at E13.5 (Figure 3A). Quantification of postnatal germ cells indicated that both IR doses used were sufficient to cause a reduction in germ cells at birth in both sexes (Figure 3B, C).

**Figure 3.**
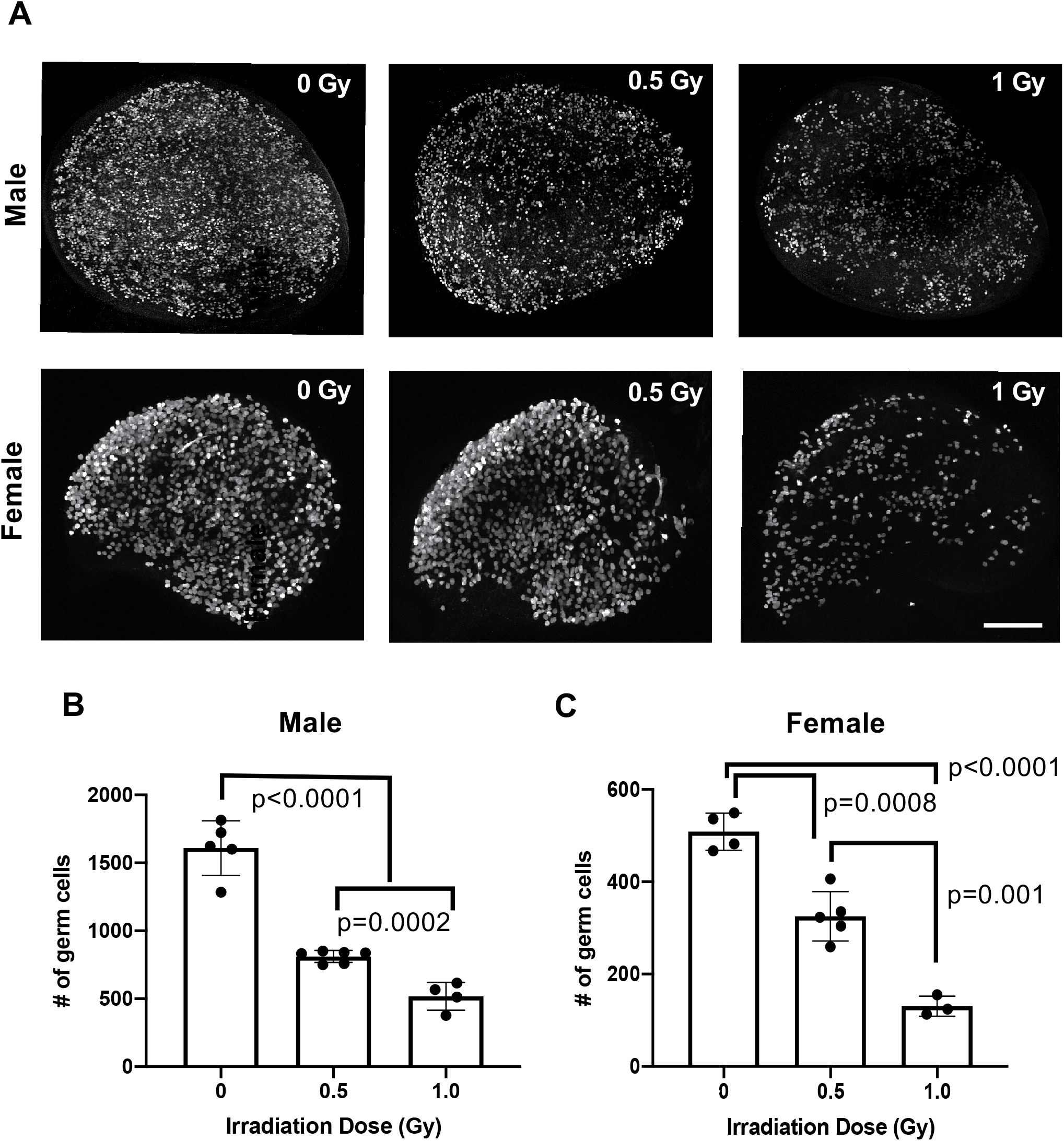
Acute radiation-induced DNA damage during fetal development leads to a reduction in germ cells at birth. (A) Whole mount maximum intensity immunofluorescence images of P0 testes and ovaries labeled with germ cell marker Tra98/GCNA in which the irradiation dose noted corresponds to the amount of radiation exposure the samples experienced *in utero* at E13.5. Scale bar=200 μM. (B) Germ cell quantification of P0 testes at the doses shown in A. Number of germ cells quantified from the 3 centermost sections of each sample is graphed; p-values derived from unpaired Student’s t-test statistical comparisons. (C) Germ cell quantification of P0 ovaries at the doses shown in A and quantified as in B; p-values derived from unpaired Student’s t-test statistical comparisons.

## Discussion

Here, we validate and demonstrate the functionality of a mouse transgenic reporter line designed to aid the study of DNA damage induction and responses in embryonic germ cells. The reporter, like the parental vector designed for generic expression in cultured cells (Dimitrova et al., 2008), was responsive to both the dose of DNA damage and time after DNA damage exposure. While numerous techniques can provide a static read-out of DNA damage, the dynamic nature of reporter allows for the examination of DNA repair activity in real time and under various different contexts. Additionally, the sensitivity of the reporter to low doses of DNA damage is conducive to further studies aimed at understanding the unique sensitivity of primordial germ cells to DNA double strand breaks.

## Materials and Methods

All graphing and statistical analyses in the study were performed using GraphPad Prism8. Statistical analyses were performed using an unpaired Student’s t-test unless otherwise noted.

### Vector construction and generation of Tg(Oct4ΔPE-53BP1mCherry)1Jcs mice

The Oct4ΔPE-53BP1mCherry construct was produced by replacing the GFP portion of the Oct4ΔPE-GFP plasmid (Addgene plasmid #52382) with a 53BP1-mCherry fragment (Dimitrova et al., 2008) (Addgene plasmid #19835). The GFP fragment was removed via *Afe*I and *Fse*I restriction enzyme digestion of the Oct4ΔPE-GFP plasmid. Re-introduction of an ~200bp conserved portion of the *Oct4* promoter adjacent to the excised GFP sequence was performed via PCR amplification (see Supplementary Table 1 for a list of primers used). During this step, a *Pac*I restriction site was also added. Next, the 53BP1-mCherry reporter was PCR amplified with primers designed to add *Pac*I and *Fse*I restriction sites to the 5’ and 3’ ends, then ligated into the recipient Oct4ΔPE plasmid lacking GFP to make the final construct for pronuclear injection.

Prior to pronuclear injection, the construct was linearized and gel purified to remove plasmid backbone. Following gel purification, the linearized Oct4ΔPE-53BP1mCherry reporter was injected at a concentration of 1.5 ng/ul into FVBxB62J F1 hybrid embryos and transgene positive animals were genotype as described below. The use of mice in this study was approved by Cornell’s Institutional Animal Care and Use Committee.

### Genotyping strategy

Crude lysates for PCR were prepared as described (Truett et al., 2000). Genotyping primers are provided in Supplementary Table 1. PCR conditions were as follows: initial denaturation at 95°C for 4 minutes, then 30 cycles of 95°C for 30 seconds, 62.5°C for 50 seconds, 72°C for 40 seconds, and a final elongation at 72°C for 5 minutes. For identification of transgene positive animals, the presence of a 131 bp PCR product was assessed on an agarose gel.

### Fertility assessments

Transgene positive and negative mice (at 8 weeks old) were housed with C57BL/6J wild type animals of the opposite sex. Litter sizes were determined by counting pups on the day of birth.

To quantify sperm, cauda epididymides from 8 week old mice were minced in 2 mL of 1X PBS and then incubated for 15 minutes at 37°C to allow spermatozoa to swim out. Sperm were then diluted 1:5 in 1X PBS and counted with a hemocytometer.

### Histology

Testes were collected from 8 week old males, fixed in Bouin’s overnight, washed with 70% ethanol for 24 hours and then embedded in paraffin. Testes were sectioned at 6 μm and stained with hematoxylin and eosin (H&E).

### Irradiation exposure

Mice were placed in a ^137^cesium irradiator with a rotating turntable and exposed to the doses of radiation specified.

### Whole mount staining

Whole mount staining was conducted as described (Rinaldi, Bloom, & Schimenti, 2018). Genital ridges were stained using primary antibody rabbit anti-DDX4/MVH (1:500, ab13840; Abcam) and anti-rabbit Alexa Fluor 488 secondary antibody (1:1000). P0 ovaries and testes were labeled with rat anti-Tra98/GCNA (1:500, ab82527; Abcam) and anti-rat Alexa Fluor 488 secondary antibody (1:1000). Samples were imaged on an upright Zeiss LSM880 confocal microscope using an either 10× NA 0.45 water immersion C-Apochromat objective (for the P0 ovaries and testes) or a 40× NA 1.1 water immersion C-Apochromat objective (for the genital ridges).

### Foci quantification

Foci quantification was performed using Arivis Vision4D 3.1.1 analysis software. The following parameters were used and applied to all images: puncta diameter=3 μm (threshold=5), nuclei diameter 14 μm (threshold=10; split sensitivity 63%), volume without holes 200 μm^3^ to 10000 μm^3^.

### Germ cell quantification

Germ cells from the 3 medial optical sections of each P0 ovary and testis collected were quantified as described in Rinaldi *et al.* 2018.

## Acknowledgments

This work was supported by National Institutes of Health grants S10-OD018516 (to Cornell’s Imaging Facility), R01HD082568 to J.C.S., and an Institutional training grant T32HD057854 that supported J.C.B. The authors would like thank R. Munroe and C. Abratte of Cornell’s Stem Cell and Transgenic Core Facility, and R. Williams for her assistance with microscopy and image analysis training.

